# Persisters are primed for CRISPR-Cas adaptation

**DOI:** 10.1101/2025.04.01.646530

**Authors:** Jesper Juel Mauritzen, Anne Sofie Wajn, Ida Friberg Hitz, Nina Molin Høyland-Kroghsbo

**Author notes:** These authors contributed equally (Jesper Juel Mauritzen, Anne Sofie Wajn). Correspondence: Nina Molin Høyland-Kroghsbo.

## Abstract

In the evolutionary battle between bacteria and mobile genetic elements, such as bacteriophage viruses and plasmids, bacteria have developed intricate defense systems. Among these, the CRISPR-Cas system has been extensively studied and harnessed as a revolutionary gene editing tool. However, while the biochemical process by which this microbial immune system acquires genetic CRISPR memory and immunity against invaders has been comprehensively examined, fundamental questions about the bacterial physiological state underlying *how* and *when* CRISPR memory is formed have only been partially explored. Naïve CRISPR adaptation is generally rare, but occurs more frequently when bacteria are challenged with replication-deficient phages. In such scenarios, bacteria are not under immediate threat and have ample time to adapt to the phage DNA, without risking cell death. Accordingly, slow growth caused by low temperatures, low aeration, or bacteriostatic antibiotics promotes CRISPR adaptation, possibly by allowing the Cas complexes more time to adapt before being outpaced. Persister cells are dormant antibiotic-tolerant subpopulations of cells with limited metabolic activity. When a mobile genetic element invades a persister cell, its replication is halted until the host cell resumes growth, providing an ideal opportunity for CRISPR adaptation. Here, we show that transiently dormant *Escherichia coli* persister cells acquire CRISPR immunity 10-fold more frequently than the general bacterial population. Thus, persister cells, in addition to being notoriously antibiotic tolerant, are primed for CRISPR-Cas adaptation and may be in a state of heightened immune capacity and evolution, securing the survival of the population.

## Introduction

Phage therapy, the use of bacteriophage viruses to treat bacterial infections, has been studied for over a century (1). Although the discovery of antibiotics reduced its use in the West, recent advancements highlight its potential (2, 3). However, its effectiveness is still limited by gaps in our understanding of bacterial defenses and stress responses. Bacteria have evolved a variety of anti-phage defenses, including the Clustered Regularly Interspaced Short Palindromic Repeats (CRISPR) and CRISPR-associated (Cas) genes (4). CRISPR-Cas systems serve as an adaptive and heritable immune system against foreign genetic elements, such as plasmids and phages (5). CRISPR-Cas operate through three main steps: adaptation, expression, and interference (5). Adaptation involves the acquisition of new spacers within the CRISPR array, creating a genetic memory of prior infections. Cas1 and Cas2 are responsible for the adaptation step (6). Expression includes the transcription of *cas* genes and CRISPR arrays, followed by the processing of pre-CRISPR RNA into mature crRNAs. Interference is the crRNA-guided destruction of target DNA executed by Cas nucleases.

Naïve adaptation towards targets not recognized by the existing crRNA repertoire is rare (7, 8) but detectable, especially in response to replication-deficient phages (9). In such scenarios, bacteria have ample time to adapt to the phage DNA without risking cell death. Slow bacterial growth, induced by factors such as low temperatures, oxygen limitation (10), and bacteriostatic antibiotics (11), also promotes CRISPR adaptation. This suggests that dormant cells, including persister cells, may be particularly prone to CRISPR adaptation. Persister cells are transiently antibiotic-tolerant dormant subpopulations that are genetically identical to the antibiotic-sensitive population (12). When a phage infects a persister cell, its replication is halted until the host cell resumes growth (13), providing an ideal opportunity for CRISPR adaptation. In this study, we explore if dormant cells, such as persister cells, are particularly prone to CRISPR adaptation.

## Results

Here we studied the type I-E CRISPR-Cas system of *Escherichia coli* MG1655 as a model system to explore whether persister cells are particularly prone to naïve CRISPR-adaptation. In *E. coli*, the *cas* genes are repressed by the global regulator H-NS under laboratory conditions (14). Therefore, we used an *E. coli* K-12 MG1655 CRISPR-adaptation system that had been previously generated (15). The strain was constructed as follows: a ΔCRISPR-*cas* strain was equipped with a minimal CRISPR array (two CRISPR repeats and a spacer from the *E. coli* MG1655 CRISPR 1 array) and carrying inducible *cas1-2* on plasmid pCas1-2 (16). Using this strain, we challenged overnight cultures with the antibiotic ciprofloxacin to select for persister cells or with Milli-Q as a general population control. The cell population collapsed upon ciprofloxacin treatment as expected (Fig. 1A). After recovering ciprofloxacin-treated cells by cultivating them without ciprofloxacin, the progeny cells were re-exposed to ciprofloxacin, which again lead to population collapse (Fig. 1B). This indicates that most of the population were not antibiotic resistant due to mutation, but rather antibiotic tolerant persister cells, as expected.

**Figure 1.**
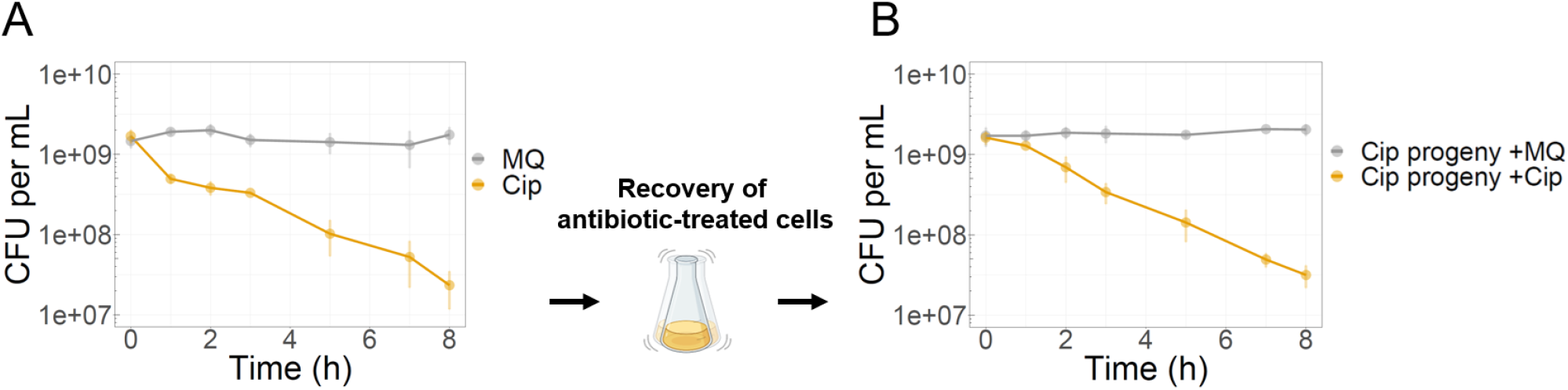
Biphasic killing curve distinguishes persistence from resistance. **A)** The general bacterial population die rapidly upon antibiotic treatment. Overnight cultures of *E. coli* MG1655 Δ*cas* ΔCRISPR *yfp*-CAR^CR-1^ pCas1+2 were cultured in LB with 50 µg mL^−1^ streptomycin, 0.1 mM IPTG and 0.2 % arabinose. Subsequently, the cultures were treated with either Milli-Q or ciprofloxacin (10 µg mL^−1^) for 8 h, washed in PBS, and plated to count colony forming units. Additionally, the washed ciprofloxacin-treated cells were resuspended in PBS, diluted to 1*10^6^ CFU mL^−1^, and grown to 8*10^8^ CFU mL^−1^ in the absence of antibiotics and inducers. **B)** Subsequently, the cultures were again challenged with either Milli-Q or ciprofloxacin, revealing that the majority of the antibiotic treatment survivors were not resistant by mutation. Error bars represent standard deviations of biological triplicates.

After the cells received their initial 8 h treatment with Milli-Q or ciprofloxacin (Fig. 1A), they were washed to remove residual antibiotics, back-diluted to a cell density of 1*10^6^ viable CFU ml^−1^, and grown to 8*10^8^ CFU ml^−1^ to achieve a cell density. We then used amplicon sequencing to quantify CRISPR spacer acquisition and found that CRISPR adaptation is 10-fold more frequent in the transiently dormant *E. coli* persister cells compared to the general bacterial population (Fig. 2A). Thus, persister cells are primed for CRISPR-Cas adaptation and may be in a state of heightened immune capacity and evolution. We also observed that, when adjusting for genome size, acquired spacers mapped to plasmid pCas1+2 at a rate 400-fold higher than to the *E. coli* chromosome (Fig 2B). Since pCas1+2 uses the CloDF13 origin of replication (16), which maintains about 10 copies per cell (17), this suggests that CRISPR adapts preferentially to plasmids by ~40-fold, supporting previous findings (18). Ultimately, persisters not only secure population survival against antibiotics (19) but are also in a heightened immune state to potentially fight off phage infections, likely limiting the success of phage-based therapies.

**Figure 2.**
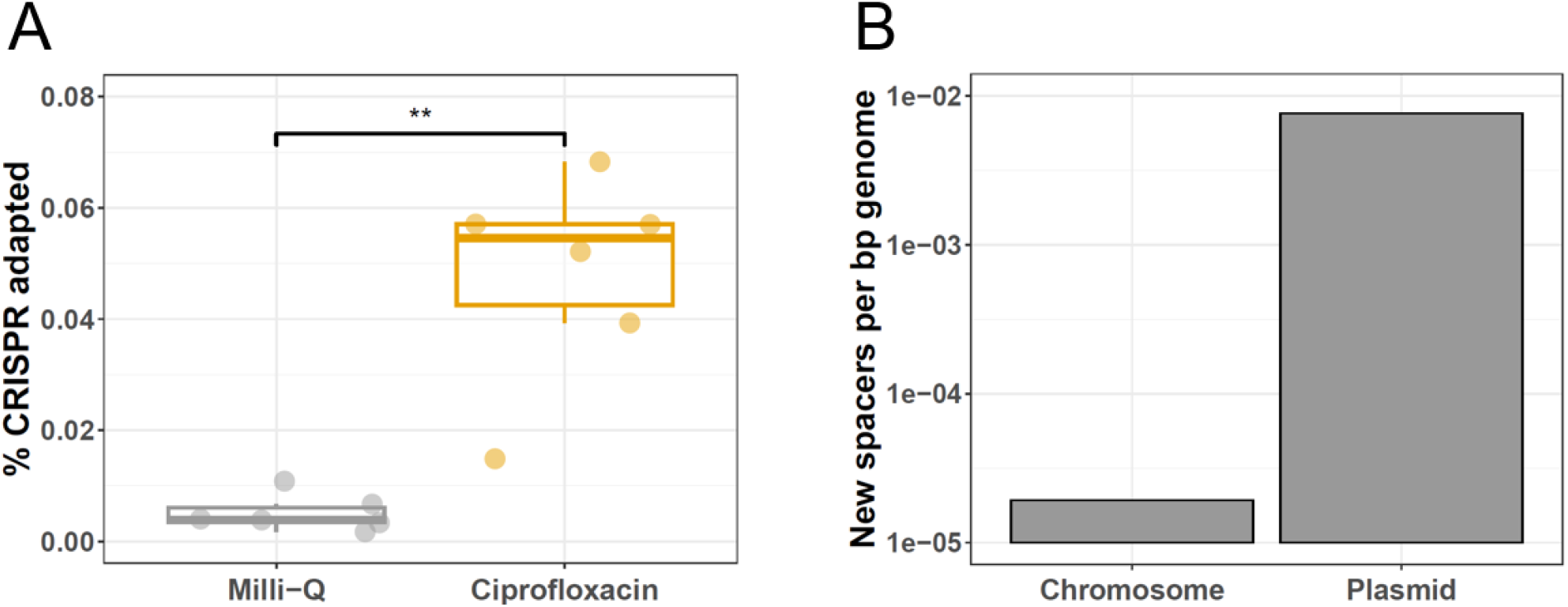
CRISPR adaptation is 10-fold more frequent in persister cells. **A)** Percent CRISPR adaptation levels of general *E. coli* populations and persister cells. Cells were cultured as in Fig. 1A and the minimal CRISPR arrays were PCR-amplified and amplicon sequenced. The acquisition of a new unique spacer was considered an adaptation event. The percent CRISPR adaptation was calculated by dividing new spacer acquisitions by the sum of adapted and non-adapted CRISPR arrays. **B)** The unique new spacers from all adaptation events were simultaneously mapped to the *E. coli* chromosome and the pCas1+2 plasmid. The number of spacers mapping to the chromosome and plasmid was normalized to genome size by dividing the new spacers by the respective genome sizes. ** represent statistical significance (unpaired *t*-test; p-value = 0.0022).

## Discussion

In this study, we have uncovered a previously overlooked phage defense mechanism where dormant persister bacteria develop CRISPR immunity 10-fold more frequently than a control population. Upon resuming growth, these adapted cells are genetically equipped to fend off infecting foreign DNA. Slow growth enhances CRISPR-Cas activity (10, 11) likely by providing additional time to adapt and dismantle invading genetic elements before they can replicate and outpace the defense mechanisms. While stalled growth increases CRISPR-Cas adaptation (Fig. 2A), our findings indicate that persister cells, cultivated for the same duration as the general population, are inherently in a state of heightened capacity for CRISPR adaptation. Ultimately, slow or stalled growth, such as in biofilm cells (20), may lead to higher CRISPR-Cas activity, making these potential environmental reservoirs for CRISPR adaptation.

Intriguingly, phage infection triggers the PQS quorum sensing stress response in *Pseudomonas aeruginosa* (21, 22), which induces persister formation (23). Similarly, viral infection in *Sulfolobus islandicus* induces dormancy (24). Consequently, this dormancy halts virus replication, allowing ample time to acquire CRISPR immunity and increasing the rate of succesfull adaptations (Fig. 2a). Furthermore, CRISPR-Cas13 activity also induces dormancy (25), suggesting that CRISPR-Cas systems may be inherently designed to slow or stall bacterial growth for optimal efficacy (Fig. 2a). Beyond CRISPR-Cas, persister cells can use a toxin/anti-toxin system to inhibit phage replication and restriction-modification systems to cleave the phage DNA during dormancy (26), indicating that multiple bacterial anti-phage defenses may be particularly potent during the persister state.

CRISPR adaptation prefers plasmid DNA over chromosomal DNA due to the higher density of Chi sites on the bacterial chromosome, limiting spacer acquisition from self-DNA (18). This bias is further influenced by the RecBCD complex, which favors plasmid DNA due to its higher number of replication forks (27). These stalled replication forks can lead to double-strand breaks that expose DNA ends, serving as templates for CRISPR adaptation (27).

Understanding immunity regulation in transiently antibiotic-resistant cells is vital for treating multidrug-resistant pathogens. Antibiotics can inadvertently select for slow-growing or dormant cells that enhance CRISPR immunity, complicating the use of phage-based therapies. This underscores the importance of strategic approaches, such as combining antibiotics with phage therapy and quorum sensing inhibitors, to synergistically eliminate pathogens, limit persister formation, and reduce the population prone to acquiring CRISPR-based immunity.

## Materials and Methods

### Bacteria and plasmids

*E. coli* K-12 strain MG1655 ΔCRISPR Δ*cas yfp*-CAR^CR1^ pCas1+2 was grown at 37 °C with shaking in Lysogeny broth (LB) or on LB agar plates solidified with 15 g agar L^−1^. To maintain plasmids, streptomycin 50 µg mL^−1^ was used. For induction of *cas1-2* expression, 0.1 mM IPTG and 0.2 % L-arabinose were added. The strains and plasmids used in this study are listed in supplementary Table S1 and S2.

### High-throughput CRISPR adaptation assay

Overnight cultures of *E. coli* strain MG1655 ΔCRISPR Δ*cas yfp*-CAR^CR1^ pCas1+2 were prepared in LB with 50 µg mL^−1^ streptomycin, 0.1 mM IPTG and 0.2 % arabinose. Subsequently, the cultures were split in two and treated with 10 µg mL^−1^ ciprofloxacin or Milli-Q as control and incubated for 8 h. Every timepoint, the cultures were centrifuged at 10,000 *g* for 2 min, washed twice in PBS, resuspended in PBS, and plated to enumerate colony forming units (CFUs). Additionally, after 8 h of incubation, the washed cells treated with Milli-Q or ciprofloxacin were diluted 1:1,000 and 1:10 in LB, respectively, to a cell density of 1*10^6^ CFU mL^−1^ and grown to 8*10^9^ CFU ml^−1^. Then, the cells were pelleted by centrifuging at 10,000 *g* for 2 min and DNA was extracted using the New England Biolabs Monarch HMW DNA Extraction Kit for Cells & Blood kit. The minimal CRISPR arrays were PCR-amplified using the Q5 polymerase with the primers listed in Table S2 and purified using the Macherey-Nagel Nuleospin™ PCR Clean-up Kit. Amplicons were sequenced by Eurofins, yielding a minimum of ~55,000 merged paired-end reads per sample. R packages ShortRead v. 1.52.0 (28) and BioStrings v. 2.62.0 (29) were used to identify repeat sequences allowing for 2 bp mismatches and counting unadapted CRISPR arrays as containing 2 repeats. For CRISPR arrays containing 3 repeats, we observed spacer1 duplicates and samples with multiple similar spacer sequences mapping to the same region and differing by only a few bp, despite the samples overall having relatively few potential acquired spacers (1-39). We deemed this biologically unlikely and attributed it to PCR biases. Therefore, newly acquired spacers were filtered to remove spacer1 duplicates and trimmed to retain only unique sequences, defining a unique sequence as having at least 4 bp difference from others. The unique spacer sequences were counted as adapted CRISPR arrays. To quantify the adaptation preference for chromosome or plasmid, the spacers were mapped simultaneously to the *E. coli* chromosome and the pCas1-2 plasmid sequences using the Geneious algorithm with the highest sensitivity and visualized using Geneious Prime v. 2022.1.1

## Statistical analysis

Unpaired *t*-test was performed using R package ggpubr v. 0.6.0 (30).

## Author contributions

A.S.W., J.J.M, and I.F.H. performed experiments. J.J.M. and N.M.H-K analyzed the data, provided supervision, and wrote the manuscript. N.M.H-K conceived the overall project idea and acquired funding.

## Acknowledgements

We thank Magnus Lundgren, Uppsala University, for kindly sharing the *E. coli* CRISPR adaptation reporter strain. We thank the Independent Research Fund Denmark, for grant 1054-00099B to N.M.H-K.

