## Supplemental material for "Persisters are primed for CRISPR-Cas adaptation"

### Supporting material

**Table S1. Bacterial strains and plasmids used in this work**

| Strain and plasmid | Description | Source |
|---|---|---|
| MG1655 ΔCRISPR Δ*cas* *yfp*-CAR$^{CR1}$ | *E. coli* strain lacking CRISPR-cas, but carrying a minimal CRISPR array | (15) |
| pCas1-2 | pCDF-1b backbone, Streptomycin resistance gene, *cas1-2* gene expression induced by arabinose and IPTG | (16) |

**Table S2. Primers used in this work.** The sequencing adaptors are highlighted in italics

| Description | Sequence 5'-3' | Source |
|---|---|---|
| *yfp* -CARCR-1 Reverse with adaptor | *ACACTCTTTCCCTACACGACGCTCTTCCGATCT*TGTTACATTAAGGTTGGTGGGTTG | This work. |
| *yfp* -CARCR-1 Forward with adaptor | *GACTGGAGTTCAGACGTGTGCTCTTCCGATCT*TGACCATGATTACGCCAAGC | (15) |
